# Phosphorylation energy and nonlinear kinetics as key determinants for G2/M transition in fission yeast cell cycle

**DOI:** 10.1101/084400

**Authors:** Teng Wang, Jian Zhao, Qi Ouyang, Hong Qian, Yu V. Fu, Fangting Li

**Affiliations:** School of Physics, Center for Quantitative Biology, Peking University, Beijing 100871, China; State Key Laboratory of Microbial Resources, Institute Of Microbiology, Chinese Academy of Sciences, Beijing 100101, China; Department of Applied Mathematics, University of Washington, Seattle, Washington 98195, U.S.A.

**Author notes:** These authors contributed equally to this work (Teng Wang, Jian Zhao).

## Abstract

The living cell is an open nonequilibrium biochemical system, where ATP hydrolysis serves as the energy source for a wide range of intracellular processes including the assurance for decision-making. In the fission yeast cell cycle, the transition from G2 phase to M phase is triggered by the activation of Cdc13/Cdc2 and Cdc25, and the deactivation of Wee1. Each of these three events involves a phosphorylation-dephosphorylation (PdP) cycle, and together they form a regulatory circuit with feedback loops. Almost all quantitative models for cellular networks in the past have invalid thermodynamics due to the assumption of irreversible enzyme kinetics. We constructed a thermodynamically realistic kinetic model of the G2/M circuit, and show that the phosphorylation energy (Δ*G*), which is determined by the cellular ATP/ADP ratio, critically controls the dynamics and the bistable nature of Cdc2 activation. Using fission yeast nucleoplasmic extract (YNPE), we are able to experimentally verify our model prediction that increased Δ*G*, being synergistic to the accumulation of Cdc13, drives the activation of Cdc2. Furthermore, Cdc2 activation exhibits bistability and hysteresis in response to changes in phosphorylation energy. These findings suggest that adequate maintenance of phosphorylation energy ensures the bistability and robustness of the activation of Cdc2 in the G2/M transition. Free energy might play a widespread role in biological decision-making processes, connecting thermodynamics with information processing in biology.

---

E. Schrödinger first suggested that living organisms require negative entropy flux to create and maintain order (*1*). From a thermodynamic perspective, the nonequilibrium nature and free energy input are indispensable for living organisms (*2,3*). Beyond the traditional view that free energy is consumed to carry out processes like biosynthesis, ionic pumping, or mechanical movements (*2*), the recent phosphorylation energy hypothesis rationalizes that free energy is a necessary component of nonlinear E. biochemical functions. Theoretical studies have shown that free energy is a decisive factor to endow biochemical reactions with ultrasensitivity, bistability or oscillation characters, and thus allows molecular players to carry out information processing and decision-making in living cells (*3–9*). Without adequate free energy input, no matter how well-designed a biochemical network is, it will not be biologically functional.

Phase transition in the cell cycle is a particularly well-characterized biological decision-making process (*10–12*). In the fission yeast, *Schizosaccharomyces pombe*, transition from G2 to M phase is triggered by the activation of cyclin-dependent kinase (CDK), a mechanism that is highly conserved from yeast to humans (*10,13,14*). Yeast CDK, Cdc2, forms a kinase complex with Cdc13, a B-type cyclin. The Cdc13/Cdc2 complex is activated by Cdc25 and inhibited by Wee1 (*14-16*) (Fig. 1A). Among these three components, there exists a positive feedback and a double-negative feedback: active Cdc13/Cdc2 complex activates the phosphatase activity of Cdc25 and inactivates the kinase activity of Wee1, both through phosphorylation on multiple sites (*17–20*). All three phosphorylation-dephosphorylation (PdP) cycles are actively driven by the free energy generated from ATP hydrolysis (*14,19,20*). Inside a living cell, the free energy generated from the hydrolysis of one mole of ATP, also called phosphorylation energy, is defined as Δ*G* =*RT* ln*γ*, where *γ*=[ *ATP*]/*K_eq_*[ *ADP*][*Pi*]) and *K_eq_* is the equilibrium constant for ATP synthesis. Under physiological conditions **γ** and Δ*G* are approximately 1010 and 57.05 kJ/mol, respectively. Δ*G* is connected with the chemical reaction fluxes and the dissipated heat of the living cell. Given that free energy Δ*G* quantifies how far a living cell is away from a thermochemical equilibrium state, a positive Δ*G* state from inanimate matter (*2-6*). distinguishes living state from inanimate matter (2–6).

**Fig. 1.**
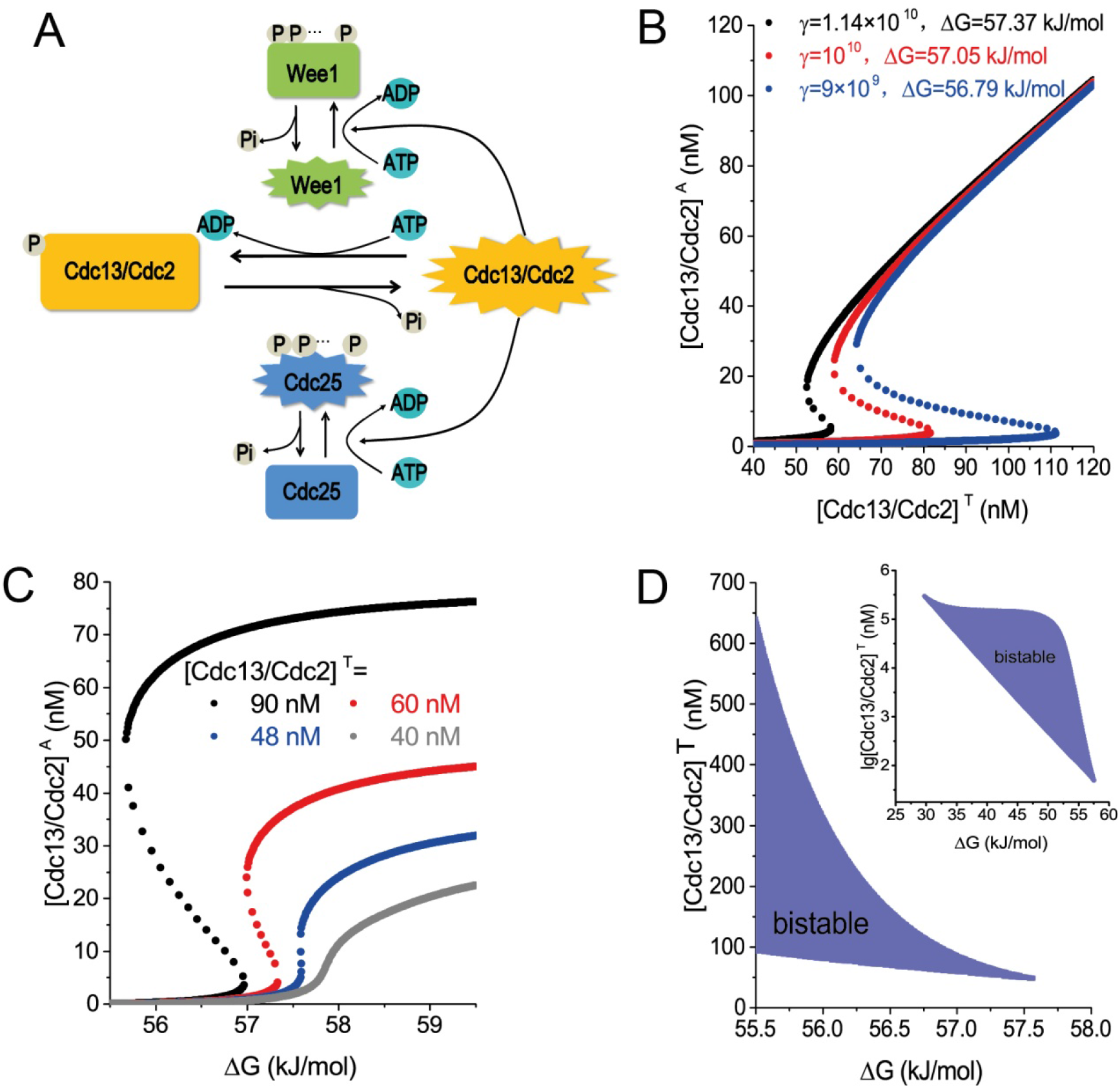
Regulatory circuit of the fission yeast G2/M transition and theoretical model predicting hysteresis in Cdc2 activation as a function of both cyclin and phosphorylation energy. (**A**) The reciprocal positive and double-negative feedback loops in the Cdc13/Cdc2-Wee1-Cdc25 circuit. The PdP cycle of Cdc2/Cdc13 is at the center. Cdc2/Cdc13 is inactive when its Tyr15 is phosphorylated via active Wee1, and this inactivation is reversed when dephosphorylation occurs due to active Cdc25. In turn, Cdc2/Cdc13 phosphorylates Cdc25 at multiple sites to activate Cdc25, and phosphorylates Wee1 at multiple sites to inactivate Wee1. (**B**) and (**C**) Modeling analysis predicts that the activity of Cdc13/Cdc2 exhibits bistability and hysteresis in response to both total cyclin ([Cdc13/Cdc2]^T^) and phosphorylation energy (ΔG). (**D**) The bistable region of Cdc13/Cdc2 activity in the G2/M system presented as functions of both Δ*G* and total cyclin. A trajectory passing through the upper boundary of the bistable region corresponds with a completion of the G2/M transition. A vertical trajectory represents the Cdc13-driving G2/M transition, and a horizontal trajectory represents a transition driven by Δ*G*.

It has been widely accepted that the strong nonlinearity generated by feedbacks results in the bistability of the system (*10,21,22*). Cdc2 activation driven by cyclin B accumulation has been experimentally demonstrated to be bistable in oocyte extracts (*21*). Thus, it is important to further quantify how the intracellular nonequilibrium environment and free energy influence cellular decision-making processes and phase transitions, especially the activation of CDK in the fission yeast G2/M transition.

To address this fundamental question, we first developed a theoretical analysis of the activation of CDK in the fission yeast G2/M transition. The existing mathematical descriptions for these biochemical networks, which are based on the assumption of irreversible PdP cycles, are thermodynamically invalid and have no role for- Δ*G* (*12,23*). In our thermodynamically realistic kinetic model, the activation-inactivation of Cdc13/Cdc2 is represented by reversible elementary kinetic steps, and the regulation of Wee1 and Cdc25 by Cdc13/Cdc2 are assumed as networks which consist of multiple PdP cycles (*19,20,24*). With some simplifications and details in section 1, 2, and 3 of supplementary text, the model gives prediction on the concentration of active Cdc13/Cdc2, [Cdc13/Cdc2]^A^ as a function of both total concentration [Cdc13/Cdc2]^T^ and Δ*G*.

The steady-state level of [Cdc13/Cdc2]^A^ can be obtained as:

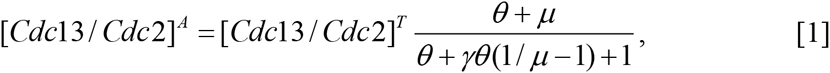

where 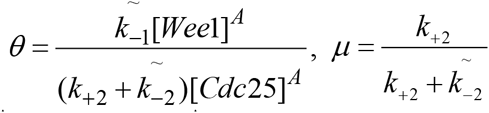, and 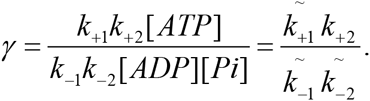 [*Wee1*]^A^ and [*Cdc25*]^A^ represent the concentrations of active Wee1 and Cdc25, respectively. *k*_+1_, *k*_-1_, *k*_+2_ and *k*_-2_ are the rate constants, and 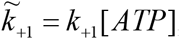, 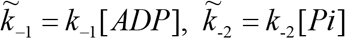. The phosphorylation energy level Δ*G* can be expressed as 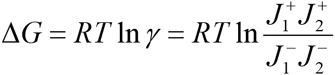 where *J*_1_^+^, *J*_2_^+^, *J*_1_^-^, and *J*_2_^-^ are the chemical fluxes generated from the activation and in activation of Cdc13/Cdc2:

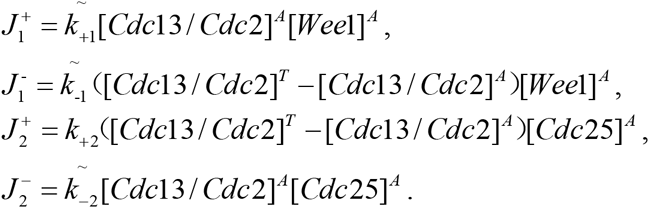

At steady state of active 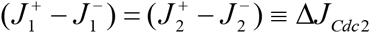. The quantity 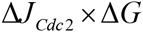 represents the heat dissipation rate in the central Cdc13/Cdc2 activation PdP cycle. We make an analogy between the biochemical circuits with the electric circuits; the phosphorylation energy Δ*G* corresponds to the voltage, Δ*J* corresponds to the current, and Δ*J* × Δ*G* corresponds to the electric power.

The G2/M transition contains multiple PdP cycles including Cdc13/Cdc2, Wee1 and Cdc25, and then we obtained the following results. Phosphorylation energy Δ*G*, the ‘voltage’ of G2/M system, governs the activities of Cdc13/Cdc2, Wee1 and Cdc25. In particularly when G2/M system reaches equilibrium state, where Δ*G*=0, *γ*=1 and Δ*J*=0, the concentration of active Cdc13/Cdc2 becomes *μ*·[Cdc13/Cdc2]^T^, which is independent of both [Wee1]^A^ and [Cdc25]^A^. Therefore, only when the system is at nonequilibrium state with Δ*G*>0 that the nonlinear activation of Cdc13/Cdc2 through Wee1 and Cdc25 could work. The key factor that governs the activation of Cdc13/Cdc2 is free energy Δ*G* or the ratio of [ATP]/([ADP][Pi]) instead of [ATP] (or [ADP]) itself.

We calculated the activation of Cdc2 stimulated by increasing cyclin B ([Cdc13/Cdc2]^T^) under various Δ*G* levels shown in Fig. 1B. We used [Cdc13/Cdc2]^T^=60 nM for the fission yeast G2 state (*25*). The lower branch, with low Cdc13/Cdc2 activity, represents the G2 state, while the upper branch, with high Cdc13/Cdc2 activity, represents the M state. The red line in Fig. 1B presents the widely accepted quantitative behavior of the G2/M transition in physiological conditions. A yeast cell in the G2 state has low [Cdc13/Cdc2]^T^ and low Cdc13/Cdc2 activity. Once the [Cdc13/Cdc2]^T^ level is increased beyond a threshold (about 82 nM in our model), the activity of Cdc13/Cdc2 increases sharply to a level that corresponds to the M state. Furthermore, Fig. 1B shows that with higher Δ*G* levels, the threshold of [Cdc13/Cdc2]^T^ required for the activation of Cdc2 becomes lower, and vice versa. Consequently, increasing the Δ*G* level is also predicted to drive the activation of Cdc2. For example, in the G2 state when [Cdc13/Cdc2]^T^ is 60 nM, if Δ*G* is increased from 57.05 kJ/mol to 57.37 kJ/mol (which corresponds to the ATP level increasing by about 14% above normal), the threshold of [Cdc13/Cdc2]^T^ required for the activation of Cdc2 drops below 60 nM. Accordingly, the Cdc2 should have already been activated to a high level before the [Cdc13/Cdc2]^T^ concentration reaches the 60 nM threshold.

Similar to cyclin B-driving Cdc2 activation, this Δ*G*-driving activation also exhibits bistability and hysteresis, as shown in Fig. 1C. Such behavior is generic according to the catastrophe theory (Fig. 1D). In a noisy cell, this bistability could prevent the high Cdc2 activity of the M state falling to the low Cdc2 activity of the G2 state when cellular ATP/ADP levels fluctuate, thereby stabilizing Cdc2 activation during the G2/M transition.

The two-dimensional parameter space (phase diagram) in Fig. 1D illustrates how the two driving factors, [Cdc13/Cdc2]^T^ and Δ*G*, synergistically determine the activation of Cdc2. The triangular shadow represents the bistable area. From the low Cdc2 activity, trajectories passing through the upper boundary that transverse the bistable region represent the transition from G2 to M phase, and are accompanied by sharply increasing Cdc2 activity. There is a critical point for the transition between monostability and bistability (Δ*G* =57.56 kJ/mol, [Cdc13/Cdc2]^T^ =48.00 nM), where the concentration of ATP is 122.9% of the normal level. This point provides both the upper limit for Δ*G* and the lower limit for [Cdc13/Cdc2]^T^, allowing bistable switch. Within physiological range, when Δ*G* decreases, more [Cdc13/Cdc2]^T^ is required for Cdc2 activation. Conversely, when [Cdc13/Cdc2]^T^ increases, the Δ*G* threshold required for Cdc2 activation is reduced. If the maximum limit of [Cdc13/Cdc2]^T^ in fission yeast is estimated to be 400 nM, or the equivalent of about 8000 molecules in a single yeast cell (*25*), Δ*G* lower than 55.72 kJ/mol (58.5% of the normal ATP concentration) will not allow cells to enter M phase According to our model, only when Δ*G* is in the range of 55.72 kJ/mol to 57.56 kJ/mol will the G2/M transition exhibit bistable switch. Further details of our model are described in Supplementary Text and presented in figs. S1–S5.

To experimentally verify the above theoretical predictions, we employed a cell-free system derived from fission yeast nuclear extracts, which is free of mitochondrial contamination. Fission yeast nuclei were isolated from G2 phase cells and crushed by high-speed centrifuge to obtain yeast nucleoplasmic extract (YNPE) (fig. S6). In the yeast cell cycle, Cdc2 activity increases gradually during G2 phase and rises to a peak (about 6-fold higher than that of early G2 phase, fig. S7A) in the M phase (*10,13,14,21*). Thus, we quantified the Cdc2 activity to monitor the G2/M transition in our system (*14,21*). We began by adding various concentrations of exogenous Cdc13 to the early G2 phase YNPE (120 min after cells were released from synchronization) and measured the Cdc2 activity. Consistent with previous reports, the low level of Cdc2 activity rose with increasing Cdc13 concentrations (fig. S7B), suggesting that our *in vitro* YNPE system faithfully simulates the biochemical reactions involving the G2/M transition (*21*).

To test the Δ*G*-driving Cdc2 activation predicted in Fig. 1C, we added ATP to the early G2 phase YNPE to increase the Δ*G* level. After various concentrations of ATP were added, we immediately measured both the activities of Cdc2 and the ATP/ADP concentrations at time zero (see section 4.4 in Supplementary Text for details). Given that the phosphate concentration was in excess in our system, we used lg([ATP]/[ADP]) to represent phosphorylation energy. We observed that the Cdc2 activity increased continuously to the M-phase level as the free energy level increased. As shown in Fig. 2A and B, increasing the exogenous ATP concentration from 1 μM to 200 μM, that is increasing lg([ATP]/[ADP]) from −3.38 to −0.12, led to a 6-fold increase in Cdc2 activity. Adding ATP to late-G2 YNPE (190 min after cells were released from synchronization) gave similar results, except that less ATP was required to drive Cdc2 activity to its highest level (Fig. 2C and D). These data suggest that Δ*G* functions as a driving factor for G2/M transition and might have a similar role to Cdc13 in this process.

**Fig. 2.**
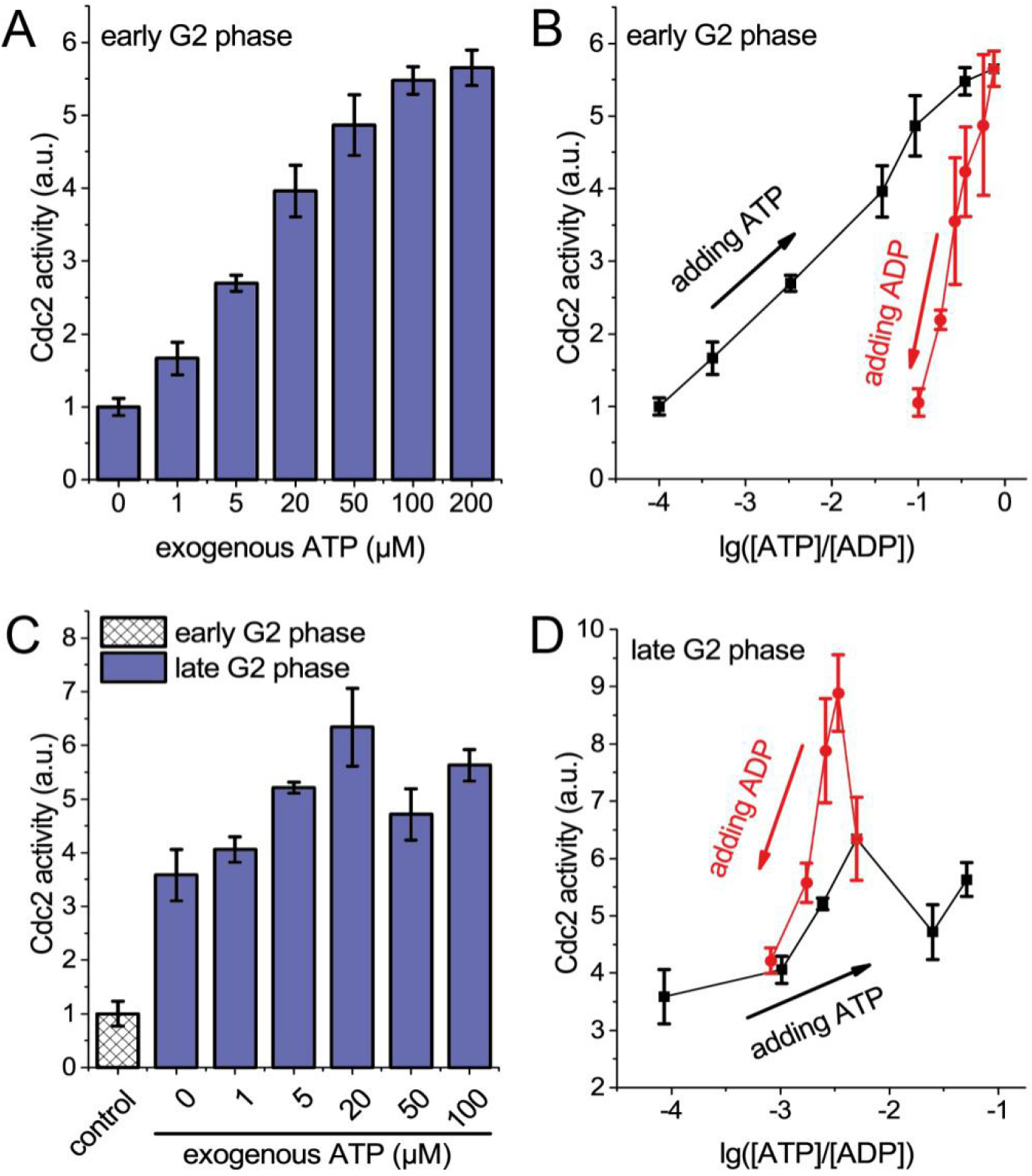
Cdc2 activation in response to Δ*G* in early and late G2 phase YNPE. (**A**) Addition of ATP to the early G2 phase YNPE (120 min after release from synchronization) increases Cdc2 activity: early G2 phase YNPE was treated with the indicated concentrations of ATP, and the Cdc2 kinase activities measured using H1 histone kinase assays. Data are normalized against Cdc2 activity in early G2 phase YNPE without exogenous ATP. (**B**) Immediately following each ATP addition, ATP and ADP concentrations were quantified: lg([ATP]/[ADP]) represents Δ*G*, and the Cdc2 activity responses to lg([ATP]/[ADP]) in early G2 phase are plotted as a black line. After 200 μM ATP was added to early G2 phase YNPE to activate Cdc2 to its M-phase level, different concentrations of ADP were added. Following each ADP treatment, the Cdc2 activity, and ATP and ADP concentrations were quantified and plotted with red line. (**C**) Addition of ATP to late G2 phase YNPE (190 min after release from synchronization) increases Cdc2 activity: as described in (**A**), using late G2 phase instead. Cdc2 activities are normalized against the Cdc2 activity in early G2 phase YNPE without exogenous ATP (indicated as control). (**D**) The Cdc2 activity in responses to lg([ATP]/[ADP]) in late G2 phase YNPE are plotted with a black line. After 20 μM ATP was added to late G2 phase YNPE to activate Cdc2 to its highest level, additions of ADP were made to reduce Cdc2 activity. The Cdc2 activity responses to lg([ATP]/[ADP]) are plotted with a red line. Error bars represent the standard deviation (SD) of three independent replicates in (**A**) to (**D**).

Next, we investigated whether bistability and hysteresis exist when Δ*G* drives Cdc2 activation. In this set of experiments, we assume that the key determinant is the phosphorylation potential; that is, the ratio of ATP to ADP concentrations. We added exogenous ATP to YNPE to boost Cdc2 activity to its highest level, followed by the addition of various concentrations of ADP to reduce Δ*G*. It is expected that if the activation process is bistable with respect to Δ*G*, M phase Cdc2 activity will remain high at the top of the “push-back” curve; namely, the M phase Cdc2 activity will be insensitive to the increase in ADP concentration. Firstly, 200 μM ATP was added to the early G2 phase YNPE, and then increasing ADP concentrations were tested. As shown in Fig. 2B, the Cdc2 activity in the early G2 phase YNPE fell abruptly when Δ*G* decreased, suggesting that no hysteresis was generated in this process. Similar treatments were applied to the late G2 phase YNPE, which contains a higher level of Cdc13. After Cdc2 activity climbed to its highest level through the addition of exogenous ATP, Cdc2 activity in the late G2 phase YNPE remained high, even when 1.2 mM ADP was subsequently added (Fig. 2D). This ADP concentration is about four times that of endogenous ADP, and Δ*G* decreases by over 2.60 kJ/mol under these conditions. This means that even when the ATP concentration decreases to 35% of its standard physiological level, Cdc2 activity remains high and the system remains in the M state. Once the M phase has been attained in the late G2 phase YNPE, Cdc2 activity is stable and insensitive to changes of Δ*G*.

The decreased Cdc2 activity observed in the early G2 phase YNPE may be explained by the Cdc13 concentration being too low to generate hysteresis, resulting in the Δ*G*-driving trajectory of the early-G2 YNPE going entirely below the bistable region established in Fig. 1D. To test this hypothesis, we supplemented the early G2 phase YNPE with 0.29 μM Cdc13. Under these conditions, the Cdc2 activation exhibited bistability and hysteresis with decreasing Δ*G* (fig. S8). The hysteresis range is greater than 2.58 kJ/mol, which is equivalent to a range of 100% to 33.9% of normal endogenous ATP. Taken together, these findings confirm that Cdc2 activation in YNPE exhibits bistability and hysteresis with decreasing Δ*G*, provided Cdc13 levels are adequate.

Having established both Cdc13 and Δ*G* as driving factors for G2/M transition and their roles in generating a bistable switch, we next addressed the relationship between Cdc13 and Δ*G* in determining G2/M transition. In light of our prediction that high Cdc13 concentrations reduce the Δ*G* threshold for Cdc2 activation (Fig. 1), we added three different concentrations of Cdc13 to the 120-min YNPE followed by additions of different concentrations of ATP. In the presence of increased Cdc13 (0.29 μM to 1.15 μM) in the YNPE, less ATP (50 μM to 0.5 μM) was required to drive the Cdc2 activity to its highest level (Fig. 3A, C and E). In terms of phosphorylation energy, the requirement for lg([ATP]/[ADP]) to yield maximal Cdc2 activity decreased from −1.01 to −2.32 (Fig. 3B, D and F). These data are in good agreement with earlier results showing that Cdc13 and Δ*G* promote Cdc2 activation synergistically.

**Fig. 3.**
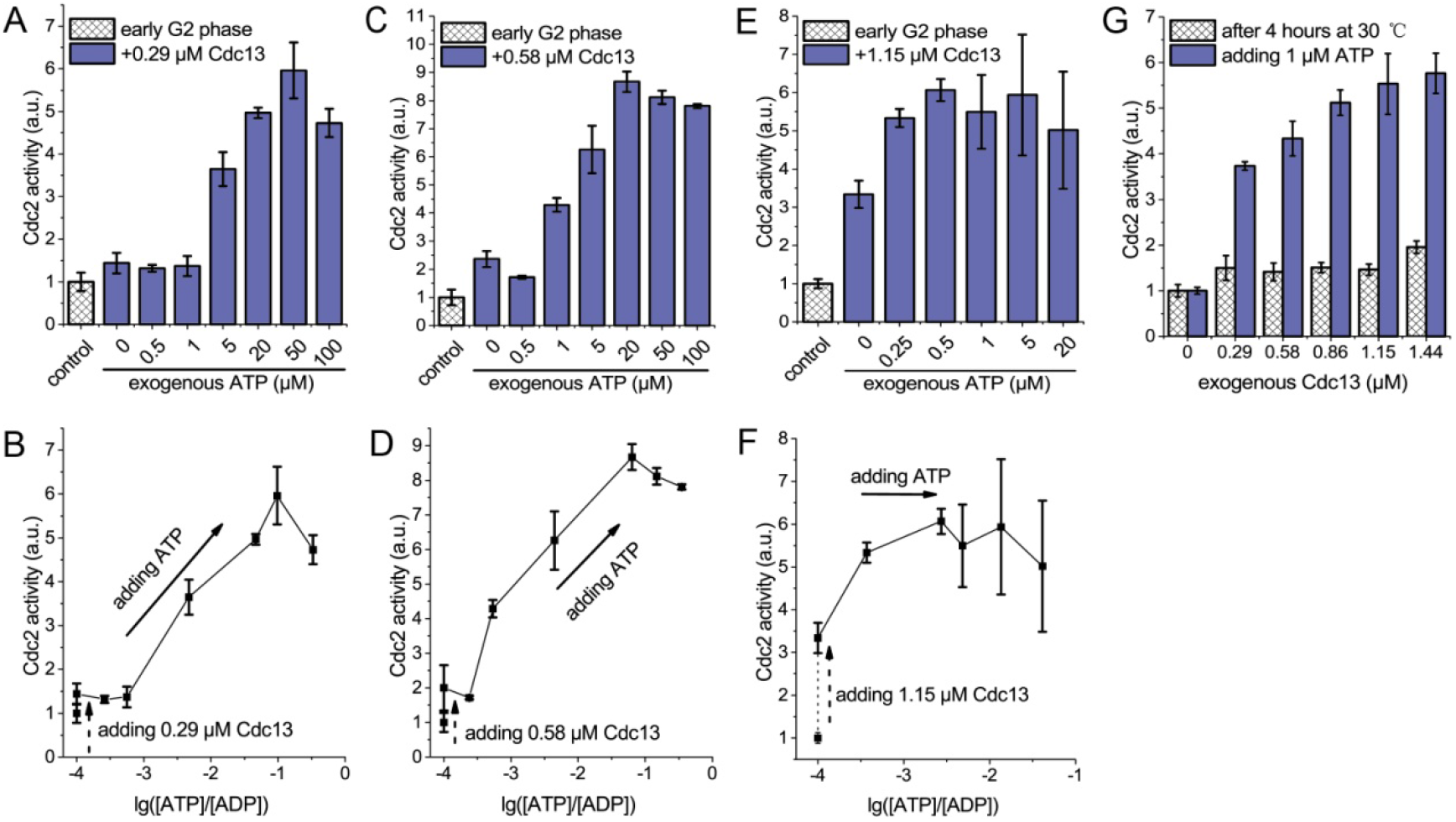
Cdc13 and phosphorylation energy synergistically trigger G2/M phase transition. (**A**)-(**F**) The 120-min YNPE was treated with 0.29, 0.58 or 1.15 μM Cdc13, and then ATP was immediately added as indicated. Cdc2 activities were measured using H1 histone kinase assays, and the ATP and ADP concentrations quantified. The Cdc2 activities in response to increasing ATP concentrations are plotted in (**A**), (**C**) and (**E**); and the Cdc2 activities in response to increasing lg([ATP]/[ADP]) are plotted in (**B**), (**D**) and (**F**). Data are normalized against the Cdc2 activity of 120-min YNPE without Cdc13 or ATP treatment (indicated as control). (**G**) The 120-min YNPE was incubated at 30°C for 4 h, and two aliquots were taken for further experiments. One aliquot was immediately treated with various concentrations of Cdc13 and the other aliquot was first treated with 1μM ATP, followed by treatments with Cdc13. Data are presented as the mean ± SD of three independent replicates.

We also explored the Cdc13-driving Cdc2 activation in extremely low Δ*G*
levels. To obtain low- Δ*G* YNPE, we incubated the early G2 phase YNPE at 30°C for 4 h, to facilitate consumption of endogenous ATP. After incubation, endogenous ATP was undetectable (data not shown) and Δ*G* was estimated to be less than 46 kJ/mol. In contrast to normal YNPE, Cdc2 in low- Δ*G* YNPE failed to be activated even following addition of 1.44 μM Cdc13 (Fig. 3G). However, if low- Δ*G* YNPE was first supplemented with 1 μM exogenous ATP (raising the Δ*G* level above 50 kJ/mol), addition of Cdc13 activated Cdc2. These results are consistent with the earlier conclusion from Fig. 1D that there exists a minimal Δ*G* threshold that permits entrance into M phase due to the maximum possible Cdc13/Cdc2 concentration in the yeast cell.

In our above experiments, ATP in YNPE was consumed and the ratio of [ATP] to [ADP] decreased over time. For the early G2 phase YNPE, we investigated the kinetics of Cdc2 activity, and ATP and ADP concentrations in the first 20 min after ATP was added (fig. S9). The curve of Cdc2 activity as a function of lg([ATP]/[ADP]) is consistent with the Δ*G*-driving Cdc2 activation curve in early G2 phase YNPE (Fig. 4A). This finding indicates that the reaction of Cdc2 activation in YNPE is fast enough that the kinetics of Cdc2 activation can indeed be treated as quasi-steady state. We also carried out theoretical simulations that supported this assumption (fig. S10, see Supplementary Text for details). Moreover, this result suggested yet another strategy to test the bistability of Cdc2 activation. Cdc2 activity in late G2 phase YNPE was measured during the first 5 min after the addition of 20 μM ATP, and an obvious bistable response of Cdc2 activity to decreasing Δ*G* was observed (Fig. 4B). A similar result was obtained for early G2 phase YNPE treated with 0.29 μM Cdc13 (Fig. 4C). Together, these data further support our conclusion above (Fig. 2) and confirm the presence of hysteresis in Δ*G*-driving Cdc2 activation.

**Fig. 4.**
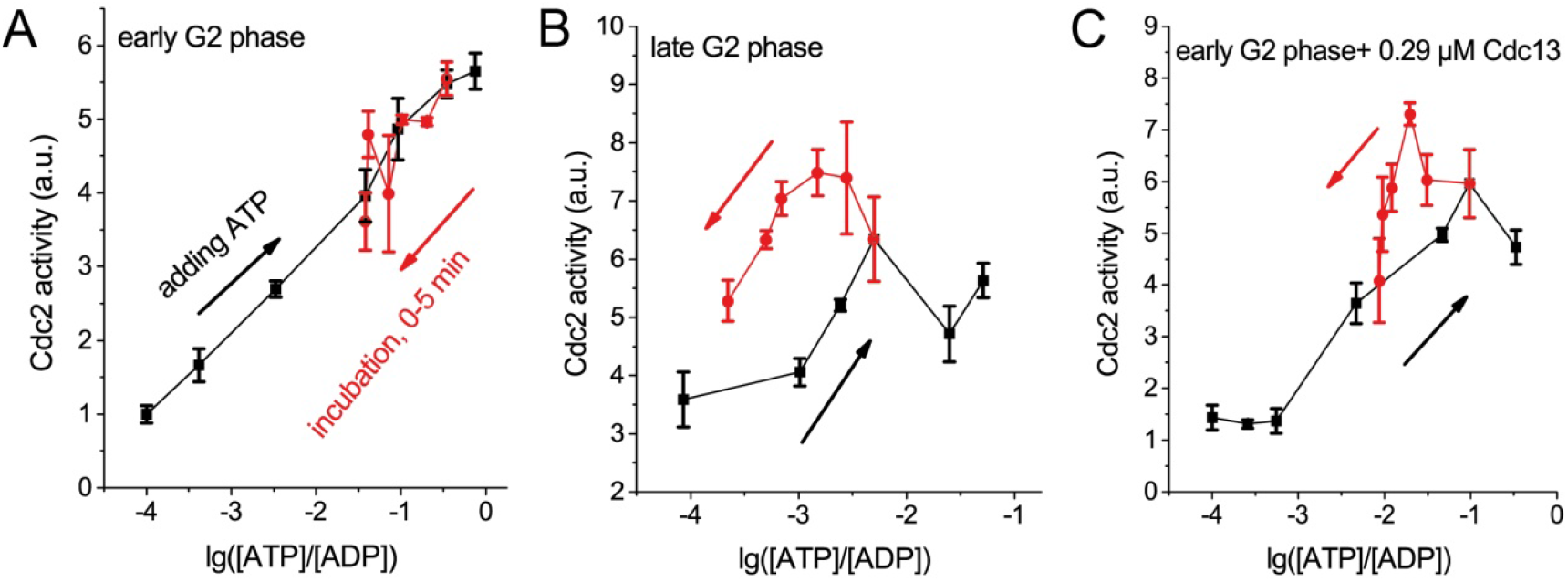
Hysteresis of Δ*G*-driving Cdc2 activation in the ATP consuming process. Pull-up curves (black) represent Cdc2 activity increasing from G2 phase state to M phase state by increasing additions of exogenous ATP. Push-back curves (red) represent declining Cdc2 activity over time when M phase state YNPE, achieved by adding exogenous ATP, was incubated at 30°C to progressively consume ATP. (**A**) 200 μM ATP was added to 120-min YNPE to increase Cdc2 activity to the M-phase level, and then incubated at 30°C for 5 min, during which the Cdc2 activity and ATP/ADP concentrations were measured every minute, The Cdc2 activity in response to lg([ATP]/[ADP]) are plotted with a red line. (**B**) 190-min YNPE was used for the same experiments described in (**A**), and 20 μM ATP was used to drive Cdc2 activity to the M-phase level. (**C**) Similar to (**A**), except that 120-min YNPE was first treated with 0.29 μM Cdc13, after 50 μM ATP was added to drive Cdc2 activity to the M-phase level. Error bars represent the SD of three independent replicates in each experiment.

From a physical stand point, a living cell is an open system that continuously exchanges chemicals with its environment in the form of both high chemical potential (cp) “food” and low-cp “waste”. A significant portion of the exchanged chemicals (for example, glucose) are used to produce ATP for intracellular *energy* (*2*), while others are used for sensing environmental *information* (for example, endocrine hormones and pheromones). The cellular decision-making processes are regulated by the free energy of ATP hydrolysis. The free energy is determined by the cellular ATP/ADP ratio or the forward/backward chemical reaction fluxes ratio, it connects with the dissipate heat and the entropy production, and quantitatively depicts the nonequilibrium status of system.

Our results strongly support this hypothesis. We showed that only when the eukaryotic yeast cell is far from equilibrium with a suitable range of free energy will the CDK activation circuit in G2/M transition exhibit the expected ultrasensitive switch with bistability and hysteresis by saddle-node bifurcation. Moreover, this saddle-node bifurcation is globally robust against cellular fluctuations of both phosphorylation energy and protein concentration. Several recent modeling studies have also revealed how the free energy and nonequilibrium govern the cellular regulatory functions and information processes (*6,9,26–29*). In this work, we directly measure the “thermodynamic force” of a living cell in the form of Δ*G* and show that it is only when it is within a specific range that the cell behaves normally, suggesting that there is a requirement for a suitable distance far from the equilibrium state. The present work clearly suggests a nontrivial interplay between regulatory circuit function, stability and nonequilibrium conditions that deserves further investigation. We also noted in Fig. 1C, through saddle-node bifurcation, that the functional circuit with suitable phosphorylation energy resides on the upper, “animated branch” of the bifurcation, while the lower, “inanimate branch” is expected to connect with the equilibrium state (Δ*G* =0). The nonlinear bistability and saddle-node bifurcations driven by Δ*G* ensure thermodynamics far from equilibrium and maintains functional robustness again perturbations. This interplay of nonequilibrium and nonlinearity allows the execution of irreversible cellular processes and possibly benefits developing new characters in further biological evolution (*10,30–32*).

Here we observe that the G2/M transition requires significant Δ*G* commitment. Accumulating evidence suggests that active Cdc2 enters mitochondria and promotes cellular ATP generation at the G2/M transition (*33,34*). Notably, in comparison with normal cells, mitochondrial respiration and glycolysis in cancer cells are altered to produce more ATP (*35,36*). Since tumor cells are in an actively dividing state, it is advantageous for them to adopt the most efficient strategy to ensure successful and ongoing cell cycles. In light of the metabolic costs and physical limitations of producing excess cyclin within cells, increasing Δ*G* to reduce the required cyclin concentration threshold seems to be an optimal alternative approach for cancer cells. Therefore, we rationalize the possibility of developing new therapies based on decreasing Δ*G* in cancer cells.

